# The formation of *Chlamydia*-containing spheres, a novel non-lytic egress pathway of the zoonotic pathogen *Chlamydia psittaci*

**DOI:** 10.1101/2023.08.28.555065

**Authors:** Jana Scholz, Gudrun Holland, Michael Laue, Sebastian Banhart, Dagmar Heuer

**Author notes:** Address correspondence to Dagmar Heuer,.

## Abstract

Egress of intracellular bacteria from host cells and cellular tissues is a critical process during the infection cycle. This egress process is essential for bacteria to spread inside the host and can influence the outcome of an infection. For the obligate intracellular Gram-negative zoonotic bacterium *Chlamydia psittaci* little is known about the mechanisms resulting in chlamydial egress from the infected epithelium. Here, we describe and characterize a novel non-lytic egress pathway of *C. psittaci* by formation of *Chlamydia*-containing spheres (CCS). CCS are spherical, low phase contrast structures surrounded by a phosphatidylserine exposing membrane with specific barrier functions. They contain infectious progeny and morphologically impaired cellular organelles. The formation of CCS shares characteristics of apoptotic cell death including a proteolytic cleavage of the peptide DEVD albeit independent of active caspase-3, an increase in the intracellular calcium concentration of infected cells, followed by blebbing of the plasma membrane and rupture of the inclusion membrane. Finally, infected blebbing cells detach and leave the monolayer thereby forming CCS. These results support that *Chlamydia psittaci* egresses the epithelial cell by a novel non-lytic egress pathway, a process beneficial for the bacterium, which might influence the outcome of the infection in organisms.

**Importance:** Host cell egress is essential for intracellular pathogens to spread within an organism and for host-to-host-transmission. Here, we describe CCS formation as a novel egress pathway for the intracellular, zoonotic bacterial pathogen *C. psittaci*. This non-lytic egress pathway is fundamentally different from previously described *Chlamydia* egress pathways. Interestingly, CCS formation shares several characteristics of apoptotic cell death. However, the sequence of proteolytic activity, followed by plasma membrane blebbing and the final detachment of a whole phosphatidylserine exposing former host cell is unique for *C. psittaci*. Thus, CCS formation represents a new egress pathway for intracellular pathogens that could possibly be linked to *C. psittaci* biology including host tropism, protection from host cell defense mechanisms or bacterial pathogenicity.

## Introduction

*Chlamydia psittaci* is the causative agent of the zoonotic disease psittacosis. Its natural hosts are birds, which can transmit *C. psittaci* to different mammal species including humans via inhalation of infectious aerosols (Knittler et al., 2014; Knittler & Sachse, 2015; Radomski et al., 2016). Human infections with *C. psittaci* cause typical pneumonia, with fever, chills, headache, coughing, and dyspnea. However, if left untreated, this infection can lead to severe disease and even death (Knittler et al., 2014; Knittler & Sachse, 2015; Radomski et al., 2016).

Like other chlamydial species, *C. psittaci* has a biphasic developmental cycle. This cycle is characterized by the presence of infectious, osmotically stable, but mostly metabolically inactive elementary bodies (EBs) and non-infectious, osmotically instable, intracellular replicating reticulate bodies (RBs) (Banhart et al., 2019; Knittler & Sachse, 2015; Radomski et al., 2016). Chlamydial infections start with attachment of an EB to the host cell plasma membrane, where it is endocytosed or phagocytosed by the host cell (Banhart et al., 2019; Elwell et al., 2016). *C. psittaci* builds its intracellular niche, the inclusion, with the help of specific chlamydial effector proteins. These proteins are secreted via type 3 secretion (Moore & Ouellette, 2014). At the same time, EBs differentiate into RBs and start to replicate (Abdelrahman et al., 2016). In addition, reorganization of cellular organelles is observed during infection, leading to fragmentation of the host cell Golgi apparatus and an increase in sphingolipid uptake into the inclusion (Christian et al., 2011; Heuer et al., 2009; Heymann et al., 2013; Koch-Edelmann et al., 2017; Rejman Lipinski et al., 2009). Interfering with ceramide to sphingomyelin conversion by chemical modified ceramide derivates dramatically decreases uptake of sphingolipids into to the inclusion and infectious progeny formation (Banhart et al., 2014). After bacterial replication and growth of the chlamydial inclusion, RBs redifferentiate into EBs. Finally, the intracellular development of *C. psittaci* is completed by host cell egress (Elwell et al., 2016; Zuck et al., 2016). For different chlamydial species, two forms of egress are described: a lytic egress and a non-lytic egress pathway called extrusion formation (Hybiske & Stephens, 2007; Zuck et al., 2016). Extrusion formation has been predominantly studied for the human pathogen *C. trachomatis*, but it is rarely investigated for *C. psittaci*. During *C. trachomatis* infections, extrusion formation depends on a cytoskeleton- and calcium-dependent packaged release process without host cell death (Hybiske & Stephens, 2007; Lutter et al., 2013; Nguyen et al., 2018). In addition, RBs can also spread between cells using tunneling nanotubes (Jahnke et al., 2022).

Egress of intracellular pathogens is of general importance for the host, as it is typically linked to tissue inflammation and induces the inflammatory response of the host organism. In comparison to other life cycle phases of intracellular pathogens, host cell egress is still understudied and thus, the identification and characterization of novel egress strategies continues (Flieger et al., 2018; Friedrich et al., 2012). In principle, egress is defined as the release of progeny to infect a new host cell. Flieger et al. (2018) distinguish between three principle egress mechanism: (1) active host cell destruction, (2) induced membrane-dependent egress without host cell destruction, and (3) induced programmed host cell death. This induced programmed host cell death can be apoptosis, pyroptosis or necroptosis (Flieger et al., 2018). During egress by active host cell destruction, the intracellular pathogens are released to the extracellular space. In addition, membrane- or cell death-dependent egress mechanisms exist, where pathogens are released into a membranous compartment to spread between cells without contact to the extracellular environment. This could protect the pathogen from extracellular immune defense mechanisms (Flieger et al., 2018; Friedrich et al., 2012). Apoptosis as egress mechanism has been reported for the Gram-positive bacterium *Mycobacterium marinum*, the Gram-negative bacterium *Salmonella enterica,* the protozoan parasite *Leishmania* spp., and the fungal pathogen *Cryptococcus neoformans* (Davis & Ramakrishnan, 2009; Flieger et al., 2018; Knodler et al., 2010). Interestingly, 25 years ago it was reported that *C. psittaci* infection causes the induction of apoptosis in late infections (Gibellini et al., 1998; Ojcius et al., 1998; Perfettini et al., 2002). However, a link of *Chlamydia*-induced apoptosis to *C. psittaci* egress was not investigated. In general, the findings about the role of different cell death pathways during chlamydial infections is still controversial and the role of apoptosis during chlamydial infections is still under investigation (He et al., 2022; Kerr et al., 2017; Matsuo et al., 2019; Sixt, 2021; Sixt et al., 2019; Waguia Kontchou et al., 2022; Weber et al., 2017).

As a hallmark of apoptosis in non-infected cells, membrane stability is influenced at different phases of the apoptotic process: At early apoptosis, phosphatidylserine is exposed to the outer leaflet of the plasma membrane and at late apoptosis, secondary necrosis can occur which includes cell membrane lysis (Lee et al., 2013; Mariño & Kroemer, 2013; Rogers et al., 2017; Zhang et al., 2018). In addition, an important regulator of apoptosis is calcium (Orrenius et al., 2003; Sukumaran et al., 2021). Early during apoptosis, calcium from the endoplasmic reticulum flows into the cytoplasm, where it triggers the release of cytochrome c from mitochondria. This induces apoptosome formation and initiates the caspase cascade (Boehning et al., 2003; Mattson & Chan, 2003). Calcium signaling has also been shown to play a role in the egress of *C. trachomatis* (Hybiske & Stephens, 2007; Nguyen et al., 2018), however, its importance for egress of *C. psittaci* is still unclear. Additionally, recent data show that *C. trachomatis* blocks store-operated calcium entry in infected host cells by recruitment of the stromal interaction molecule 1 (STIM1) and the inositol-trisphosphate receptor (Chamberlain et al., 2022).

Here, we describe a novel, non-lytic egress mechanism of *C. psittaci* that involves formation of *Chlamydia*-containing spheres (CCS). This egress pathway is observed after detection of a proteolytic activity predominately in late *C. psittaci* inclusions and cytosolic calcium concentration increases in infected cells. After calcium increase, the plasma membrane starts blebbing and the inclusion membrane ruptures. In contrast to *C. trachomatis* extrusions, CCS are formed by the detachment of the infected cells and the resulting CCS contain cellular organelles besides *Chlamydia* and expose phosphatidylserine at the surrounding membrane.

## Material and Methods

### Cell culture and infection assays

Cell culture and infection assays were performed as described previously (Koch-Edelmann et al., 2017). For further information, please see S1 Text.

### Live cell imaging of CCS

To monitor the formation of CCS, live cell imaging of either fluorescently stained *C. psittaci*-infected cells or *C. psittaci*-infected cells stably expressing GFP was performed. For more details on both methods, please see S1 Text.

### Reinfection assay

Infectious EBs were quantified as described before (Aeberhard et al., 2015). In brief, HeLa cells were infected with *C. psittaci* and incubated for 48 h. Then, supernatants were collected and centrifuged at low speed (5 min, 300 x *g*, RT) to pellet CCS and separate them from free bacteria. Subsequently, total supernatants, CCS and free bacteria were subjected to glass bead lysis followed by infection of freshly seeded HeLa cells. Cells were fixed at 24 h pi, stained for Hsp60 (mouse anti-Hsp60 (bacterial) (1:600; catalog no. ALX-804-072; Enzo Life Sciences), and numbers of infection-forming units (IFU) per mL were calculated from average inclusion counts in 10 fields of view per condition.

### Quantitative real-time PCR

Determination of bacterial genome copy numbers by quantitative real-time PCR (qRT-PCR) was performed as already described (Lienard et al., 2011) using samples from the reinfection assay.

### Isolation and staining of CCS

CCS in the supernatant of *C. psittaci*-infected cell cultures were separated by centrifugation (5 min, 300 x *g*, RT) at 48 h pi. For live staining, pellet was mixed with the indicated staining solutions. The preparation of the staining solutions and the staining procedures are described in S1 Text. For immunofluorescence staining, pellet was mixed with 4% paraformaldehyde in PBS and transferred into a poly-L-lysine coated 8 well chambered coverslip (µ-Slide 8 Well, Ibidi). After 30 min of incubation at RT, staining was continued as described for immunofluorescence assays.

### Immunofluorescence assay

For immunofluorescence assays, HeLa cells were seeded onto cover slips and infected with *C. psittaci* or left uninfected. At indicated time points, culture medium was removed and cells were fixed with 4% paraformaldehyde in PBS (30 min, RT). After washing with PBS, samples were blocked and permeabilized using 0.2% Triton X-100 and 0.2% BSA in PBS (20 min, RT). Incubation with primary antibodies (1 h, RT) was followed by washing and incubation with secondary antibodies and DAPI (25 µg ml^-1^, Sigma-Aldrich) (1 h, RT). All antibodies were diluted using 0.2% BSA in PBS. The used antibody concentrations are listed in S1 Test. All samples were analyzed at a Stellaris 8 Confocal Microscope (Leica Microsystems).

### Transmission electron microscopy

CCS in the supernatant of *C. psittaci*-infected cell cultures were separated by centrifugation (5 min, 300 x *g*, RT) at 48 h pi and the sediments were fixed with a mixture of glutaraldehyde and formaldehyde (2.5% and 1%, respectively; in 50 mM HEPES buffer). After incubation for 2 h at RT, sediments were embedded in low-melting point agarose. Samples were post fixed in osmium tetroxide (1% in distilled water) followed by block contrasting with tannic acid (0.1 % in 50 mM HEPES buffer) and uranyl acetate (2% in distilled water). Subsequently, samples were dehydrated by incubation in a graded ethanol series and embedded in epon resin. After polymerization for 48 h at 60 °C, ultrathin sections (approx. 70 nm) were prepared using an ultramicrotome (UC7; Leica) and stained with uranyl acetate (2% in distilled water, 20 min) followed by lead citrate (2 min) to increase contrast. Sections were examined with a transmission electron microscope (Tecnai 12, FEI Corp.) at 120 kV. Images were recorded using a CMOS-camera (Phurona, emsis). Overview images of entire section profiles of CCS were acquired by using the image montaging mode of the camera software. Images were aligned and stitched by using Microsoft Image Composite Editor.

### Quantification of CCS

The number of CCS in *C. psittaci*-infected cell cultures was determined under different inhibitory and medium conditions. Information about the different conditions and experimental procedures are given in S1 Text.

### Determination of proteolytic DEVD cleaving activity

To determine proteolytic DEVD cleaving activity, cells were cultured and infected in 8 well chambered coverslips (µ-Slide 8 Well, Ibidi). Proteolytic DEVD cleaving activity was determined using the Incucyte Caspase-3/7 Red Dye for Apoptosis (Sartorius) as described in the manufacturer’s protocol. For more details, see S1 Text.

### Calcium imaging

Imaging of intracellular calcium levels and distribution was performed by either Rhod 3 staining or live cell imaging of the dual calcium reporter cell line HeLa ER-LAR-Geco G-Geco (Stelzner et al., 2020). Both methods are described in more detail in S1 Text.

## Results

### Formation of *Chlamydia*-containing spheres (CCS) represent the predominant egress pathway of *C. psittaci* and is characterized by membrane blebbing and rupture of the inclusion membrane

Besides host cell invasion and intracellular replication, host cell egress is one of the essential developmental steps in the life cycle of intracellular pathogens (Flieger et al., 2018). Chlamydial egress has mainly been described for *C. trachomatis* as being either host cell lysis or non-lytic extrusion formation (Hybiske & Stephens, 2007; Zuck et al., 2016). However, little is known about egress pathways of the zoonotic pathogen *C. psittaci*. Thus, we aimed to investigate *C. psittaci* egress in greater detail. First, we monitored *C. psittaci* infected HeLa cells stably expressing eGFP by live cell microscopy starting at the end of the chlamydial developmental cycle (45 h pi). We observed the formation of spherical, low phase-contrast structures in the supernatant of the infected cell cultures that we called *Chlamydia*-containing sphere (CCS) (Figure 1 A). They were formed by a non-lytic egress pathway which started with blebbing of the host cell, followed by enlargement of the blebs and host cell detachment, thereby forming the CCS. During CCS formation, inclusions were initially intact as they were devoid of cytosolic eGFP, but then turned eGFP-positive, demonstrating rupture of the inclusion membrane during the process.

**Figure 1.**
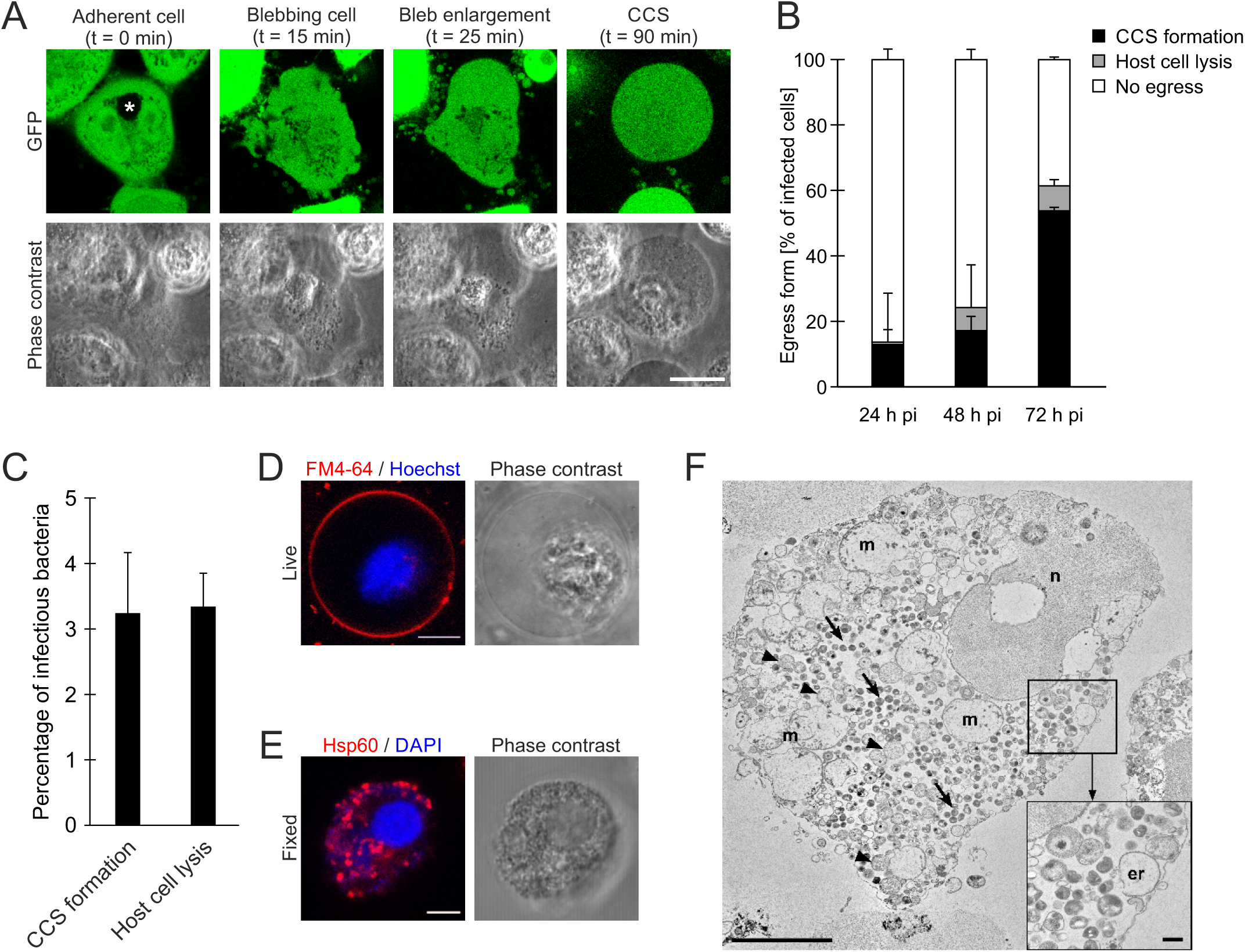
CCS represent the predominate *C. psittaci* egress characterized by lysis of the inclusion membrane and plasma membrane blebbing. (A) Time course of CCS formation. *C. psittaci*-infected HeLa cells (MOI 2) stably expressing GFP were monitored starting at 44 h pi using a CLSM equipped with a live-cell chamber. Panels show representative images of a CCS forming HeLa cell at 45 h pi. Scale bar, 20 μm; n = 2. Asterisks indicates *C. psittaci* inclusion. **(B)** Frequency of different *C. psittaci* egress pathways. *C. psittaci*-infected HeLa cells (MOI 2) were fluorescently labeled with Cell Explorer Green at 20, 44 h pi or 68 h pi and Z-stack images were acquired using a CLSM equipped with a live-cell chamber for 4 h. Within the observed infected cells, cells were categorized as having formed a CCS, being lysed or showing no signs of egress during the observation time. Egress pathways of at least 100 infected cells were analyzed. Data show mean ± SEM; n = 3. **(C)** CCS preserve infectivity of *C. psittaci.* CCS in the supernatant of *C. psittaci*-infected HeLa cells (MOI 2) were separated from free bacteria by centrifugation (300 x *g*, 5 min, RT) at 48 h pi. Infectious progeny was titrated after glass bead lysis and numbers were normalized to genome copy numbers determined by qRT-PCR. Data show mean ± SEM; n = 3. **(D)** Representative fluorescence images of a live CCS isolated of the supernatant of *C. psittaci*-infected HeLa cells (MOI 2, 48 h pi). The surrounding membrane was visualized using the membrane marker FM 4-64 and DNA was counterstained by Hoechst. Scale bar, 10 μm; n = 3. **(E)** Representative fluorescence images of a PFA-fixed CCS isolated from *C. psittaci*-infected HeLa cells (MOI 2, 48 h pi). Bacteria inside the CCS were detected using a chlamydial Hsp60 (Cy3) antibody and the DNA was counterstained using DAPI. Scale bar, 5 μm; n > 3. **(F)** Transmission electron microscopy of a thin section through a chemically fixed CCS isolated of the supernatant of *C. psittaci*-infected HeLa cells (MOI 2, 48 h pi). The CCS is almost entirely surrounded by an intact membrane and reveals many RBs (arrowheads) and EBs (arrows) besides structurally impaired cell organelles, such as mitochondria (m) or the nucleus (n). The inset shows a small region of the rim of the CCS at higher magnification with an artificially expanded profile of the rough endoplasmic reticulum (er) and bacteria at different stage of differentiation. Scale bar of the overview = 5 µm, of the inset = 0.5 µm.

Next, we aimed to quantify the newly described egress pathway of CCS formation and compare it to host cell lysis. We determined frequencies of CCS formation and host cell lysis at different timepoints after infection by counting CCS formation and cell lysis (Figure 1 B). At 20 to 24 h pi, 13% of infected host cells formed CCS, 1% of cells were lysed and 86% of the infected cells were still adherent showing no signs of bacterial egress. At 44 to 48 h pi, the percentage of CCS forming cells and lysed cells increased to 17% and 7%, respectively, while 76% of the cells were still adherent. From 68 to 72 h pi, CCS formation represented the predominant egress pathway with 54% of the cells developing CCS compared to 8% of infected cells undergoing lysis and 39% remaining intact. Most of the generated CCS were stable during 4 h observation phase. Just a small subfraction of CCS also underwent lysis subsequent to their formation, suggesting the CCS formation delays release of cellular and bacterial material including infectious EBs, pathogen-associated molecular patterns and additional danger signals into the supernatant.

As a prerequisite for successful egress, infectious chlamydial progeny needs to be released from the infected cell monolayer. Thus, we determined the percentage of infectious bacteria inside CCS and compared it to free bacteria released during host cell lysis at 48 h pi. We collected the supernatant from infected cells and separated CCS from free bacteria by differential centrifugation. Infectious progeny was titrated after glass bead lysis and counts were normalized to genome copy numbers. CCS contained approx. 3.2 ± 0.9% of infectious progeny, which was comparable to free bacteria (3.3 ± 0.5%) (Figure 1 C), showing that CCS release infectious bacteria in quantities comparable to release by host cell lysis.

*C. trachomatis* extrusions were shown to be surrounded by a double membrane in which bacteria were found, host cell nuclei were absent, and other host cell organelles were rarely present (Zuck et al., 2017). To further compare *C. psittaci* CCS with *C. trachomatis* extrusions, we analyzed CCS by live and fixed cell confocal microscopy, detecting the plasma membrane with the membrane marker FM 4-64, DNA with Hoechst or DAPI and *Chlamydiae* with an antibody against chlamydial Hsp60. We found that CCS were surrounded by a membrane (Figure 1 D), contained bacteria that were dispersed throughout the CCS as well as host cellular DNA (Figure 1 E). CCS were further characterized by thin section transmission electron microscopy (TEM) (Figure 1 F). TEM showed that CCS are limited by a more or less intact plasma membrane and that they contain *C. psittaci* EBs and fewer RBs which are dispersed throughout the CCS. The bacteria intermingle with morphologically impaired cell organelles, such as mitochondria, endoplasmic reticulum and the nucleus, and are not separated from them by an inclusion membrane (Figure 1 F).

*C. trachomatis* extrusion formation depends on different parts of the host cell cytoskeleton (Hybiske & Stephens, 2007). To investigate the requirement of the host cell cytoskeleton for CCS formation, we used the previously described inhibitors jasplakinolide, latrunculin B, nocodazole, wiskostatin, and blebbistatin as inhibitors of the actin depolymerization, the actin polymerization, the microtubules, the neural Wiskott-Aldrich syndrome protein (N-WASP), and myosin II, respectively. Using inhibitor concentrations showing effects on *C. trachomatis* egress, we were unable to determine CCS formation after wiskostatin treatment due to massive host cell death. We observed no significant difference in the number of formed CCS between DMSO treated controls (6.6 x 10^4^ CCS/cm^2^) and jasplakinolide, latrunculin B, nocodazole, and blebbistatin treated samples (5.8, 5.3, 6.4, and 6.3 x 10^4^ CCS/cm^2^, respectively) (Supplementary Figure 1), which is in contrast to *C. trachomatis* extrusion formation.

In sum, these data show that *C. psittaci* predominantly egresses by CCS formation releasing most of the infectious progeny. Moreover, these experiments demonstrate that CCS differ from *C. trachomatis* extrusions as they contain cytosolic *Chlamydia* spread through the CCS missing an intact inclusion membrane and host cellular organelles. Furthermore, CCS formation is independent of the host cell cytoskeleton. We propose that CCS are formed by a novel non-lytic egress mechanism that is distinct from other egress pathways of intracellular bacteria.

### CCS formation correlates in time with a proteolytic activity specific for a DEVD-containing substrate independent of caspase activation

We aimed to better characterize the molecular mechanisms leading to CCS formation. About twenty years ago, it has been shown that an atypic form of apoptosis gets activated during late *C. psittaci* infection, but the function of this form of apoptosis for *C. psittaci* infection remains unclear (Gibellini et al., 1998; Ojcius et al., 1998; Perfettini et al., 2002).

By live cell microscopy, we detected plasma membrane blebbing during CCS formation, which is described as a characteristic feature of apoptosis (Aoki et al., 2020; D’Arcy, 2019). Thus, we asked whether CCS formation is linked to activation of apoptosis.

The intrinsic and extrinsic pathways of apoptosis converge in caspase-3 activation (D’Arcy, 2019). We therefore analyzed the presence of cleaved caspase-3 in *C. psittaci*-infected cells at 48 h pi by immunofluorescence staining using an antibody specific for activated caspase-3. At 48 h pi, we did not detect activated caspase-3 in the cytosol of *C. psittaci*-infected host cells whereas uninfected, staurosporine-treated cells showed a clear staining of activated caspase-3 (Figure 2 A). Interestingly, using a fluorescent caspase-3/7 detection reagent, we observed proteolytic cleavage of the DEVD-containing substrate predominantly in *C. psittaci* inclusions at 48 h pi prior to CCS formation (Figure 2 B), and in CCS themselves (Figure 2 C). We further quantified this proteolytic DEVD cleavage in adherent cells. We found that it was significantly increased towards the end of the chlamydial developmental cycle from 1.4-fold intensity compared to uninfected cells at 24 h pi to 2.2- and 1.9- fold intensity compared to uninfected cells at 32 and 48 h pi, respectively (Figure 2 D)

**Figure 2.**
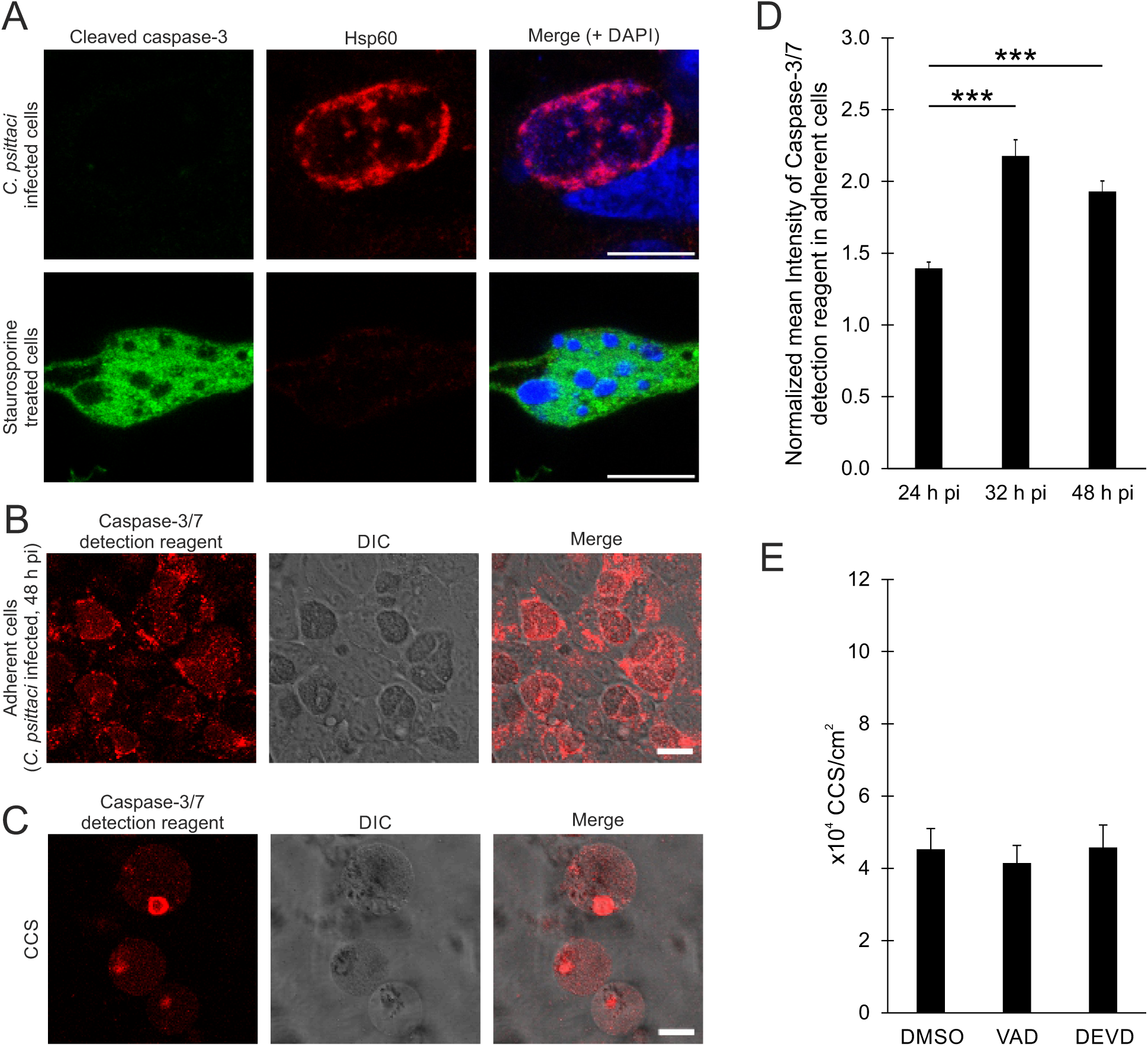
CCS formation is independent of caspases, while a DEVD-containing substrate is proteolytically cleaved during CCS formation. (A) No cleavage of caspase-3 is detectable at late *C. psittaci* infections. Representative fluorescence images of *C. psittaci*-infected HeLa cells (MOI 2, 48 h pi) and staurosporine-treated uninfected HeLa cells as positive control. Cells were fixed with PFA and *C. psittaci* and cleaved caspase-3 were detected using a mouse-anti-Hsp60 (Cy3) and a rabbit-anti-cleaved caspase-3 (AF488) antibody, respectively. The DNA was counterstained using DAPI. Scale bar, 10 μm; n = 2. (B, C) *C. psittaci* infection leads to activation of the caspase detection reagent by proteolytic cleavage of a DEVD containing substrate. HeLa cells were infected with *C. psittaci* (MOI 2) and at 40 h pi, the cells were stained with Incucyte Caspase-3/7 Dye for Apoptosis. Representative images of the infected, adherent cells **(B)** and of CCS **(C)** in the supernatant at 48 h pi are shown. Scale bar, 20 μm; n = 3. **(D)** Quantification of the proteolytic activity during CCS formation. *C. psittaci*-infected HeLa cells (MOI 2) were stained with Incucyte Caspase-3/7 Dye for Apoptosis at 16 h pi, 24 h pi or 40 h pi for 8 h. Z-stack images were acquired at a CLSM. Data were normalized to mean fluorescence intensity of uninfected cells at respective time points. Data show mean ± SEM; n = 3; *p < 0.05; **p < 0.01; ***p < 0.005 (Student’s t-test). **(E)** CCS formation is not affected by caspase inhibition. *C. psittaci-* infected HeLa cells (MOI 2) were treated with 2 µM Z-VAD-FMK, 2 µM Z-DEVD-FMK or DMSO (negative control) at 44 h pi. CCS in the supernatant were quantified at 48 h pi. Data show mean ± SEM; n = 3.

Next, we asked if the inhibition of caspase-3 influences CCS formation. After treatment with Z-VAD-FMK as pan-caspase inhibitor and Z-DEVD-FMK as a specific caspase-3 inhibitor from 44 to 48 h pi, we observed no significant difference in the number of CCS between DMSO treated controls (4.5 x 10^4^ CCS/cm^2^) and Z-VAD-FMK and Z-DEVD-FMK treated samples (4.1 and 4.6 x 10^4^ CCS/cm^2^, respectively) (Figure 2 E) indicating that CCS formation is independent of caspase activity.

These data indicate that CCS formation is independent of a caspase activity and that a DEVD-containing substrate is cleaved within *C. psittaci* inclusions prior CCS formation.

### The surrounding CCS membrane expose phosphatidylserine and retains selective permeability barrier function

During early apoptosis, phosphatidylserine is exposed to the outer leaflet of the plasma membrane (Lee et al., 2013; Mariño & Kroemer, 2013). In addition, at late stages of apoptosis secondary necrosis can be induced, leading to the loss of plasma membrane integrity (Rogers et al., 2017; Zhang et al., 2018). Thus, we focused on the surrounding membrane of CCS. To test for apoptosis characteristics late during *C. psittaci* infections when CCS formation is observed, we stained CCS with annexin V as phosphatidylserine marker and SYTOX Green as marker of plasma membrane integrity. In the CCS fraction, we observed a membrane staining with annexin V around CCS (Figure 3 A), but no SYTOX Green staining within CCS (Figure 3 B). We also observed nuclei of dead, lysed cells in the CCS fractions that were positive for SYTOX Green staining. Quantification revealed that all stained CSS were annexin V-positive whereas only 8% of CCS were SYTOX-positive (Figure 3 C, D).

**Figure 3.**
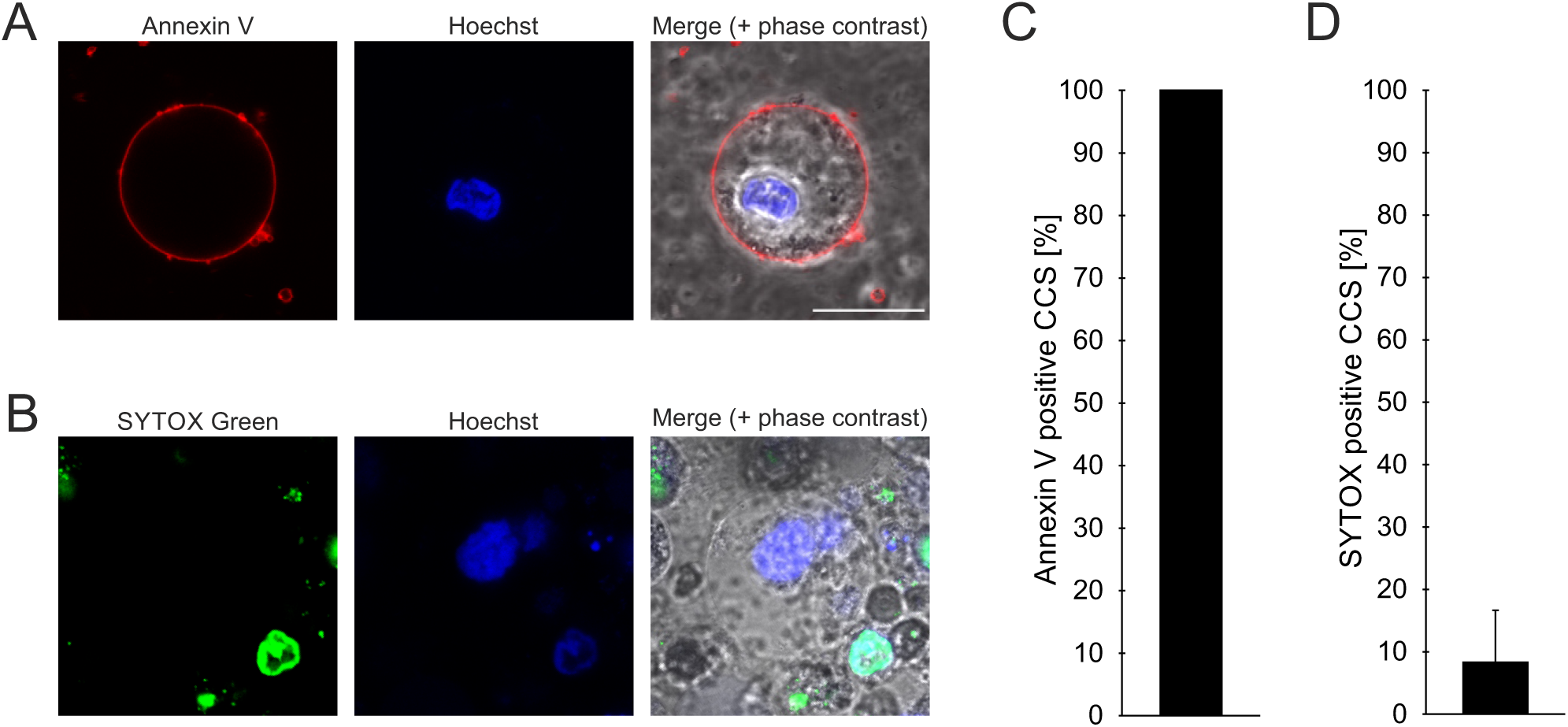
CCS present phosphatidylserine at their membrane surface and retain membrane integrity. (A, B) Representative images of a CCS positive for Annexin V (Alexa Fluor 568) **(A),** but not for SYTOX Green **(B)**. CCS were collected from the supernatant of *C. psittaci*-infected HeLa cells (MOI 2, 48 h pi), DNA was counterstained with Hoechst. Scale bar, 20 μm; n = 3. **(C, D)** Quantification of phosphatidylserine exposure **(C)** and membrane integrity **(D)** of CCS. CCS were collected from the supernatant of *C. psittaci*-infected HeLa cells (MOI 2, 48 h pi) and either PS was visualized using Annexin V (Alexa Fluor 568) **(C)** or the nucleic acids of CCS with damaged or compromised membranes were stained with SYTOX Green **(D)**. DNA was counterstained using Hoechst. The number of DNA-containing CCS surrounded by an Annexin V staining **(C)** or CCS with Hoechst and SYTOX Green stained DNA **(D)** was determined and normalized to the total amount of CCS with Hoechst stained DNA. Data show mean ± SEM; n = 3.

These data show that membranes of CCS remain intact and in contrast to *C. trachomatis* extrusions, expose phosphatidylserine on the outer leaflet. Besides membrane blebbing, the phosphatidylserine exposure on the outer leaflet is another indicator for contribution of apoptosis to CCS formation.

### CCS formation is sensitive to changes in the extracellular calcium concentration and cytosolic calcium concentration increases prior to CCS formation

Another well characterized regulator of apoptosis is calcium (Mattson & Chan, 2003; Pinton et al., 2008; Sukumaran et al., 2021). Furthermore, calcium has been described to regulate the egress of *C. trachomatis* (Hybiske & Stephens, 2007; Nguyen et al., 2018), thus we investigated the role of calcium on the formation of CCS.

We first tested if changes in extracellular concentration of calcium alters CCS formation. When we cultivated *C. psittaci*-infected cells with calcium chloride concentrations ranging from 0 to 400 mg L^-1^ from 44 to 48 h pi, we observed a dose-dependent increase in CCS formation until 200 mg L^-1^ calcium chloride (from 4.2 to 21.7 x 10^4^ CCS/cm^2^), while CCS numbers decreased at 400 mg L^-1^ of calcium chloride to 12.5 x 10^4^ CCS/cm^2^ (Figure 4 A).

**Figure 4.**
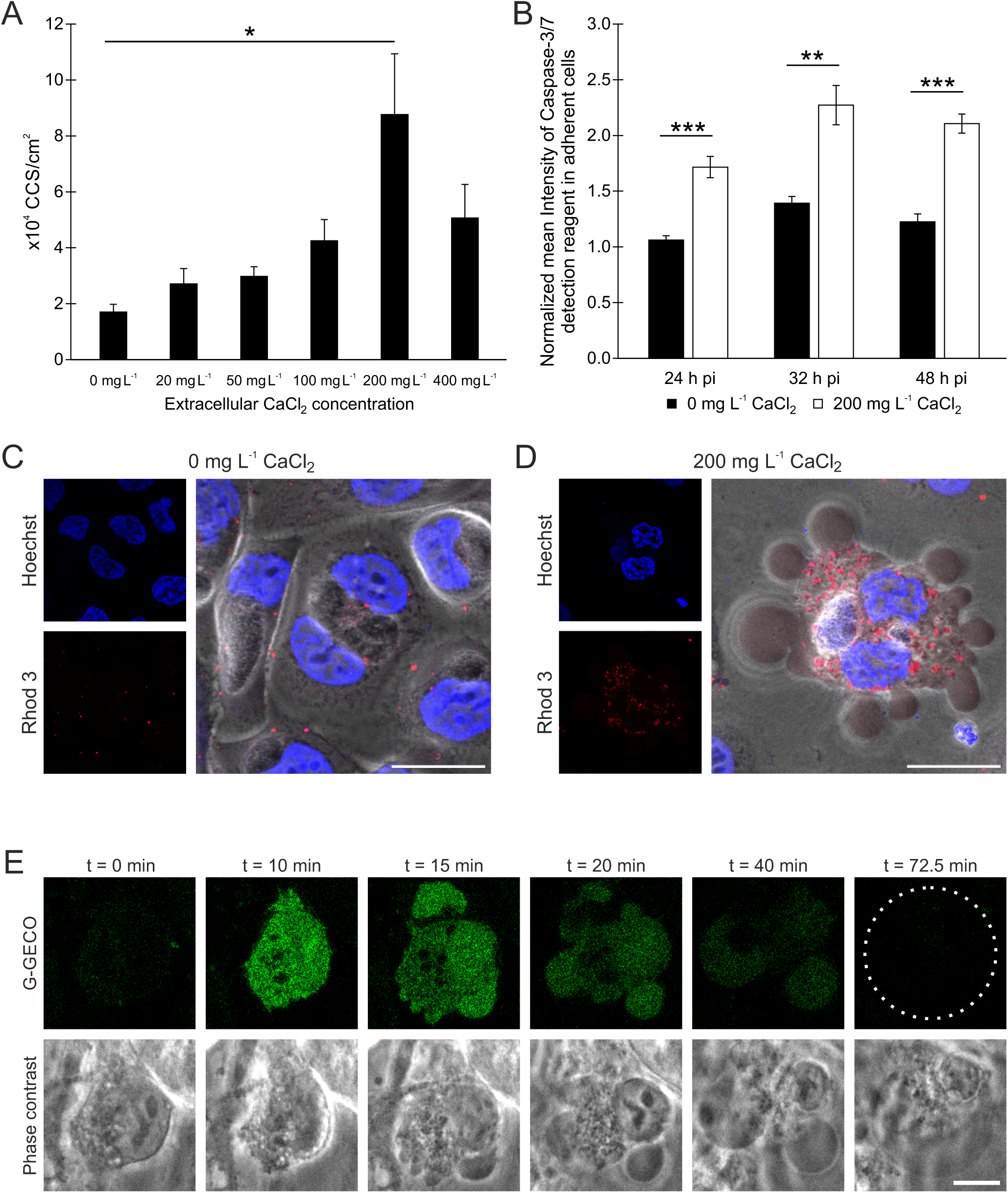
CCS formation is calcium dependent. (A) CCS formation depends on the extracellular calcium concentration. HeLa cells were infected with *C. psittaci* (MOI 2). At 44 h pi, culture medium was replaced by serum-free medium supplemented with the indicated concentrations of calcium chloride. CCS in the supernatant were visually quantified at 48 h pi. Data show mean ± SEM; n = 4; *p < 0.05; **p < 0.01; ***p < 0.005 (Student’s t-test). **(B)** Proteolytic DEVD cleaving-activity in *C. psittaci*-infected HeLa cells depends on the extracellular calcium concentration. Culture medium of *C. psittaci*-infected HeLa cells (MOI 2) was replaced by serum-free medium supplemented with indicated concentrations of calcium chloride at 20 h pi, 28 h pi and 44 h pi and cells were stained with Incucyte Caspase-3/7 Dye for Apoptosis for 4 h. Z-stack images were acquired using a CLSM. Data were normalized to mean fluorescence intensity of uninfected cells without calcium chloride at respective time points. Data show mean ± SEM; n = 3; *p < 0.05; **p < 0.01; ***p < 0.005 (Student’s t-test). **(C, D, E)** Intracellular calcium concentration increases during CCS formation. **(C, D)** HeLa wildtype cells were infected with *C. psittaci* (MOI 2). At 44 h pi, medium was replaced by serum-free medium supplemented with 0 mg L^-1^ calcium chloride **(C)** or 200 mg L^-1^ calcium chloride **(D)**. At 46 h pi, cells were labeled with the calcium sensor Rhod-3 for 1 h. At 48 h pi, DNA was counterstained using Hoechst and images were acquired. Representative images of intact cells (0 mg L^-1^ L CaCl_2_) and CCS formation (200 mg L^-1^ CaCl_2_) are shown. Scale bar, 25 μm; n = 2. **(E)** Dual calcium reporter HeLa ER-LAR-Geco G-Geco cells were infected with *C. psittaci* (MOI 2). At 44 h pi, cells were monitored at a CLSM equipped with a live-cell chamber. Panels show representative images of a CCS forming cell. Dashed line indicates CCS. Scale bar, 10 μm; n = 3.

Next, we addressed if extracellular calcium concentration affected cleavage of the DEVD-containing substrate late in *C. psittaci* infected cells. For this, we compared the DEVD cleaving activity at 200 mg L^-1^ calcium chloride containing conditions to calcium-free conditions using the caspase-3/7 detection reagent. At all examined time points (24, 32 and 48 h pi), we observed a decrease in the intensity from 1.7- to 1.1-fold, 2.3- to 1.4-fold, and 2.1- to 1.2-fold, respectively (Figure 4 B).

Subsequently, we analyzed the intracellular calcium concentration during late stages of *C. psittaci* infection using the membrane permanent cytosolic calcium sensor Rhod-3 at calcium-free and 200 mg L^-1^ calcium chloride containing conditions. At calcium-free conditions, we did not detect an increased cytosolic calcium concentration in *C. psittaci*-infected cells at 48 h pi (Figure 5 C), while at 200 mg L^-1^ calcium chloride, we observed *C. psittaci*-infected cells with blebbing membranes and an increased cytosolic calcium concentration (Figure 4 D).

**Figure 5.**
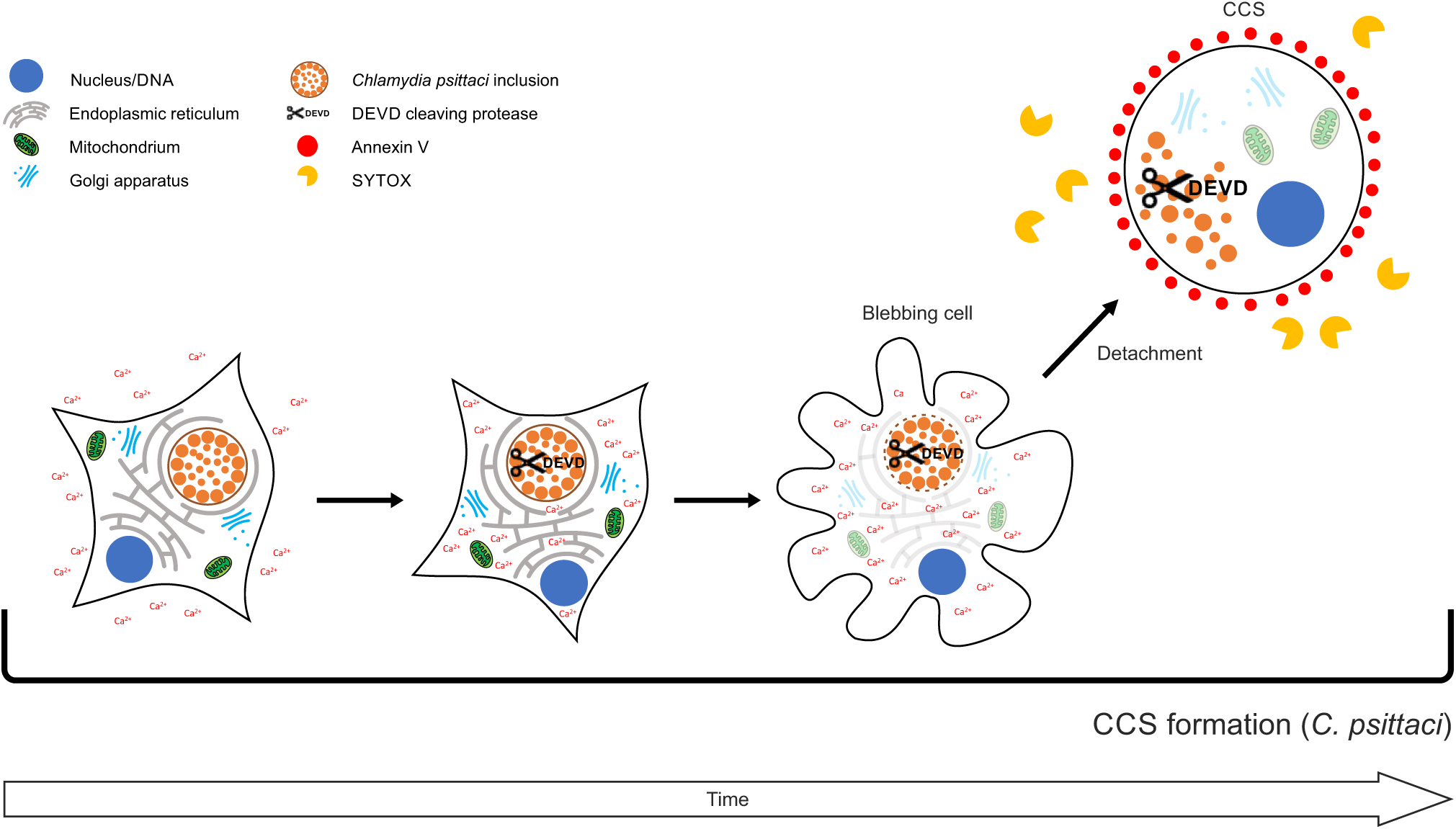
Graphical model of *C. psittaci* CCS formation. See discussion for details.

To determine the chronological order of membrane blebbing and calcium influx, we microscopically visualized intracellular calcium levels during CCS formation using a genetically encoded calcium sensor (Stelzner et al., 2020). Using HeLa cells that stably express G-Geco, an increase in intracellular calcium concentration was detected during CCS formation before initiation of the CCS characteristic membrane blebbing. This increased calcium concentration is subsequently slowly reduced during the CCS formation, lead to a CCS with again basal calcium levels (Figure 4 E).

These data show that the cytosolic calcium concentration is increased prior to CCS formation affecting the cleavage of a DEVD-containing substrate and CCS formation. In sum, our data support that calcium is an important regulator in *Chlamydia* biology especially in controlling egress pathways of *C. psittaci* and *C. trachomatis*.

## Discussion

Egress of infectious bacteria from an infected host cell or tissue is essential for intracellular bacteria to finish their infection cycle and spread within a host or transmit to a new host. Differences in egress pathways have been linked to specific aspects of the biology of pathogens including transmission, host tropism and pathogenicity (Flieger et al., 2018; Kerr et al., 2017; Spera et al., 2023; Traven & Naderer, 2014). Thus, understanding egress strategies used by different pathogens appear as important as the understanding of adhesion and invasion processes to gain deeper insights into the biology of a specific pathogen. Interestingly, for the zoonotic bacterial pathogen *C. psittaci* little is known about egress mechanisms.

In this study, we demonstrate that *C. psittaci* uses a novel non-lytic pathway to egress from an epithelial cancer cell culture. This pathway uses the formation of host cell derived *Chlamydia*-containing spheres (CCS). CCS are found in the supernatant of infected cell monolayers and are limited by a single membrane that originated from the host cell plasma membrane. The surrounding membrane maintains selective membrane barrier functions and exposes phosphatidylserine on the outer leaflet. Besides infectious EBs and RBs, CCS contain host cell DNA and morphologically impaired host cell organelles, including the nucleus. CCS formation starts after detection of a caspase-3-like protease activity predominately inside inclusions, followed by an increase in cytosolic calcium concentration, lysis of the inclusion membrane, and plasma membrane blebbing. Subsequently, the infected cell detaches and leaves the monolayer resulting in CCS in the supernatant (Figure 5).

For members of the family Chlamydiaceae, egress has mainly been studied for the strict human pathogen *C. trachomatis*. It has been shown, that *C. trachomatis* can egress the infected epithelial host cell by lysis of the host cell or extrusion, a non-lytic egress pathway. Both pathways are mutually exclusive and during *C. trachomatis* infection of HeLa cell culture monolayers both pathways appear at nearly identical frequencies (Hybiske & Stephens, 2007). Interestingly, *C. psittaci* preferentially egresses the host cell by non-lytic CCS suggesting that at least in cell culture less cells would be lysed by *C. psittaci* infections compared to infections with *C. trachomatis*.

In addition to the observed differences in frequencies, CCS are morphologically distinct and their formation is mechanistically clearly distinguished from *C. trachomatis* extrusions. The increase in proteolytic activity against a DEVD-containing substrate preferentially inside *C. psittaci* inclusions before inclusion membrane rupture suggests that a currently unknown protease might be involved in inclusion membrane lysis and in initiating CCS formation. Interestingly, the involvement of a potential calcium-sensitive protease during CCS formation is further supported by our observations showing a calcium dependency of the detected protease activity and inhibition of CCS formation under calcium limitation conditions suggesting that a currently unknown protease activity affects non-lytic egress by *C. psittaci*. Previous reports showing that *C. psittaci* infection induces an atypical form of apoptosis that could not be inhibited by caspase inhibitors also hints to a role of a currently unknown protease in *C. psittaci* biology (Gibellini et al., 1998; Ojcius et al., 1998; Perfettini et al., 2002). This hypothesis is in contrast to the role of proteases in *C. trachomatis* egress. In *C. trachomatis* infections, the pan-cysteine protease inhibitor E-64 and an inhibitor of cellular calcium-dependent calpains prevented lytic egress (Hybiske & Stephens, 2007; Kerr et al., 2017). Thus, identification and characterization of the protease active during *C. psittaci* infections is important to understand the role of cellular and/ or bacterial proteases in controlling egress pathways exploited by different *Chlamydia* spp.

For *C. trachomatis* infections, calcium signaling regulates both lytic and non-lytic egress (Hybiske & Stephens, 2007; Nguyen et al., 2018). Ionized calcium is a second messenger in eukaryotic host cells and an increase in intracellular calcium concentration is a cellular danger signal that triggers the activation of cellular proteases and lipases (Orrenius et al., 2003; Sukumaran et al., 2021). Activated hydrolases can harm the membranes such as the plasma membrane and the cytoskeleton (Cai et al., 2016; Vakifahmetoglu-Norberg et al., 2013). Thus, cellular calcium homeostasis is tightly regulated by calcium specific proteins including pumps, channels and sensors (Berridge et al., 2003). SOCE is an essential pathway that regulates cellular calcium. Molecularly, the ER-localized calcium sensor STIM1 senses calcium reduction in the ER and translocates from the ER to the plasma membrane (Gudlur et al., 2018; Hogan, 2015; Venkatachalam et al., 2002). At the plasma membrane, STIM1 and Orai1 interact generating calcium influx channels inducing SOCE. Cytosolic calcium concentration increases by influx of extracellular calcium that refills the calcium in the ER subsequently (Hogan, 2015; Lunz et al., 2019). Interestingly, during *C. trachomatis* infections SOCE is impaired and recruitment of STIM1 to *C. trachomatis* inclusions has been implicated in SOCE inhibition (Chamberlain et al., 2022). CCS formation by *C. psittaci* depends on the extracellular and intracellular calcium concentration. Calcium-free medium inhibits CCS formation. In addition, we detected an increase in intracellular calcium concentration before membrane blebbing and lysis of the inclusion membrane was observed. We therefore, hypothesize that late during *C. psittaci* infections cellular calcium homeostasis is affected and results in an influx of extracellular calcium into the cytosol that triggers lysis of the inclusion membrane followed by plasma membrane blebbing which promotes CCS formation.

The inclusion represents a protective intracellular niche that supports chlamydial growth. Therefore, lysis of the inclusion membrane needs to be regulated during the chlamydial cycle of development to avoid premature inclusion membrane lysis that would activate host cell death pathways and limit the formation of infectious EB. Interfering with expression of specific *C. trachomatis* Inc-proteins by generating *C. trachomatis* mutants induced premature lysis of the inclusion membrane (Bishop & Derré, 2022; Dimond et al., 2021; Sixt et al., 2017; Weber et al., 2017). Recent studies by Bishop & Derré (2022) showed that presence of IncS stabilizes the inclusion membrane in a species dependent manner supporting previous findings using *C. trachomatis* and *C. muridarum* chimeras (Dimond et al., 2021). IncS is an inclusion membrane protein that is found at *C. trachomatis* ER-inclusion membrane contact sites and recruits the cellular calcium sensor STIM1 to these sites (Agaisse & Derré, 2015; Bishop & Derré, 2022). During the early phase of the development, functions of IncS seems to be conserved between different *Chlamydia* species whereas its role in inclusion membrane stability at later time points was species dependent (Bishop & Derré, 2022; Cortina & Derré, 2023; Dimond et al., 2021). Interestingly, depletion of STIM1 did not rescue late inclusion lysis phenotype of IncS deficient *C. trachomatis* mutants (Bishop & Derré, 2022). In contrast, depletion of STIM1 reduced extrusion formation in *C. trachomatis* (Nguyen et al., 2018). Using live cell microscopy with *C. psittaci* infected and eGFP-expressing HeLa cells, we noticed that rupture of the inclusion membrane was seen after an increase in intracellular calcium concentration – followed by CCS formation. In addition, we did not detected STIM1 at *C. psittaci* inclusions by indirect immunoflourescence microscopy (data not shown). Thus, it remains to be determined, how the inclusion membrane is lysed before CCS formation. Further experiments are needed to identify the precise role of cellular calcium signaling and the cellular calcium sensors STIM1 during infections with different *Chlamydia* species, especially in regulating egress of *C. psittaci* by CCS.

In recent years, several studies revealed that rupture of the inclusion membrane is coupled to cytotoxicity of the infected host cell but the role of distinct cell death pathways is still controversial (Bishop & Derré, 2022; Dimond et al., 2021; Kerr et al., 2017; Sixt et al., 2017; Weber et al., 2017). Interfering with expression of specific *C. trachomatis* Inc-proteins by generating *C. trachomatis* mutants induced premature lysis of the inclusion membrane and cell death by apoptosis, necrosis and aponecrosis (Bishop & Derré, 2022; Dimond et al., 2021; Sixt et al., 2017; Weber et al., 2017). Using a laser ablation strategy to artificially ruptured *C. trachomatis* inclusion membrane, Kerr et al. (2017) showed that after inclusion membrane rupture, the cells underwent necrosis independent of caspases. Interestingly, we detected the cleavage of a caspase-3 fluorogenic substrate, in both late inclusions and CCS, in the absence of active caspase-3 confirming previous observations that cellular caspase-3 is not activated during late *C. psittaci* infections (Ojcius et al., 1998; Perfettini et al., 2002). Furthermore, we observed increase of the intracellular calcium concentration, plasma membrane blebbing and exposure of phosphatidylserine at the CCS membrane surface. While these observations support a role of apoptosis in CCS formation, following observations contradict this view. CCS formation could not be inhibited by different caspase inhibitors including the pan-caspase inhibitor z-VAD-fmk and nuclear condensation was not detected prior CCS formation supporting previous findings on induction of an atypical form of apoptosis during *C. psittaci* infections (Ojcius et al., 1998; Perfettini et al., 2002).

In comparison to other intracellular pathogens, egress mechanisms similar to but not identical with CCS formation are described. An example for egress in host cellular membrane covered structures is the formation of merosomes during *Plasmodium* infections. Here, after rupture of the parasitophorous vacuole membrane, the host cell detaches and subsequently, *Plasmodium* liver stage parasites are released into vesicles that bud off from the infected host cell (Burda et al., 2017; Shears et al., 2019). In contrast to CCS formation, membrane blebbing does not occur prior to host cell detachment and phosphatidylserine is absent from the merosome (Baer et al., 2007).

Furthermore, *S. enterica*-infected epithelial cells can extrude from infected monolayers while undergoing inflammatory cell death (Co et al., 2019; Knodler et al., 2010, 2014). During this process, caspase 1, 3, and 4 are activated, leading to the formation of an epithelial cell-intrinsic noncanonical inflammasome and interleukin 18 release (Knodler et al., 2010, 2014). Similar to CCS formation, the bacteria in this extruded epithelial cell are not surrounded by a vacuolar membrane (Knodler et al., 2010). For *Listeria monocytogenes* and *Rickettsia parkeri*, the formation and extrusion of three-dimensional host cell mounds, which were replaced by the neighboring uninfected cells, were observed, which was interpreted as an innate immunity-driven process. However, this process is host cell cytoskeleton-dependent, includes NF-κΒ, but no caspase activation and occurs only in low-infected cell populations (Bastounis et al., 2021; Co et al., 2019). Interestingly, the detachment of the infected host cell during CCS formation also occurs without damaging the surrounding cell monolayer. Thus, we speculate that CCS formation could reduce the activation of the host defense system and tissue inflammation during *C. psittaci* infections. Accordingly, this suggests a link between CCS formation and pathogenicity of *C. psittaci* and should therefore be studied in the future.

Taken together, we demonstrate that *C. psittaci* predominantly egresses by the novel non-lytic mechanism of CCS formation. The sequence of caspase-independent cleavage of a DEVD-containing substrate, calcium influx, inclusion membrane lysis, plasma membrane blebbing and detachment of the whole infected host cell represents a new egress strategy for intracellular pathogens. In particular, this novel non-lytic egress mechanism is distinct from extrusion formation, the previously described non-lytic egress mechanism of the related but strict human pathogen *C. trachomatis*. Thus, we speculate that CCS formation could be linked to the specific biology of the zoonotic pathogen *C. psittaci*. Future studies in more complex cell culture models are needed to shed light on the benefits of specific egress strategies exploited by distinct chlamydial pathogens.

## Acknowledgements

We thank Andrea Martini, Madita Winkler and Henning Krüger for excellent technical assistance. The authors would like to thank Christine Suetterlin and Ming Tan (both University of California, Irvine) for critical comments on the manuscript. We would also like to thank Kathrin Stelzner and Thomas Rudel (both Universität Würzburg) for providing us with the dual calcium reporter cell line HeLa ER-LAR-Geco G-Geco. This work was financially supported by the Deutsche Forschungsgemeinschaft (www.dfg.de): SPP 2225 to Dagmar Heuer (HE 6008/4-1).

## Appendix

### Supplemental Figures

**Supplementary Figure 1.**
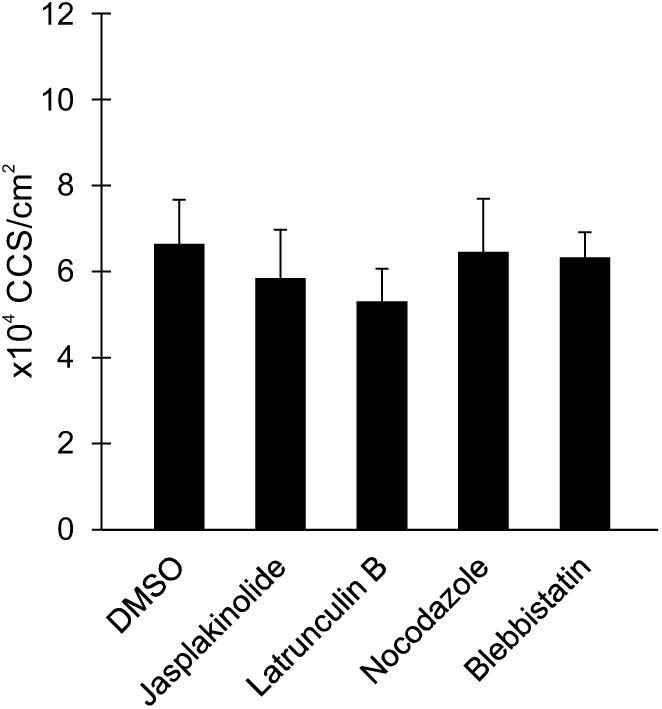
CCS are formed independently of the host cell cytoskeleton. CCS formation is not influenced by inhibitors of host cell cytoskeleton elements. HeLa cells were infected with *C. psittaci* (MOI 2) and treated with 1 µM jasplakinolide, 0.5 µM latrunculin B, 30 µM nocodazole, 50 µM blebbistatin or DMSO (negative control) at 44 h pi. CCS in the supernatant were quantified at 48 h pi. Data show mean ± SEM; n = 4.

### Supplemental Experimental Procedures

#### S1 Text

##### Cell culture and infection assays

Cell culture and infection assays were performed as described previously (Koch-Edelmann et al., 2017). Briefly, HeLa cells (ATCC CCL-2) were cultivated in RPMI 1640 medium (Gibco) supplemented with 10% fetal bovine serum (FBS, Sigma-Aldrich), 1 mM sodium pyruvate (Gibco), and 2 mM L-glutamine (Gibco) at 37°C and 5% CO_2_ and passaged every 2 to 3 days. For infection assays, subconfluent HeLa cells were washed with infection medium (Dulbecco’s modified Eagle’s medium (DMEM, Gibco) with glucose (4.5 g L^-1^) supplemented with 5% FBS, 1 mM sodium pyruvate, and 2 mM L-glutamine) and incubated with *C. psittaci* strain 02DC15 (Goellner et al., 2006) at 35°C and 5% CO_2_ using the indicated multiplicity of infection (MOI). Infected cell cultures were centrifuged (30 min, 600 x *g*, room temperature (RT)) at 30 min post infection (pi) and washed with infection medium at 2 h pi.

##### Live cell imaging of CCS

To monitor the formation of CCS, live cell imaging of either fluorescently stained *C. psittaci*-infected cells or *C. psittaci*-infected cells stably expressing GFP was performed.

For live cell labeling experiments, cell culture and infection were performed in 8 well chambered coverslips (µ-Slide 8 Well, Ibidi). *C. psittaci*-infected HeLa cells were fluorescently labeled at indicated time points using Calcein Green (AAT Bioquest) as described in the manufacturer’s protocol. In brief, culture medium was removed and 250 µL per well of Calcein Green working solution were added. After incubation (30 min, 35°C, 5% CO2), cells were washed three times with infection medium and 250 µL infection medium were added for live cell microscopy. Live cell microscopy was performed at an LSM 780 confocal laser scanning microscope (CLSM) (Carl Zeiss) equipped with a live cell chamber at 35°C. Z-stacks of 12 slices with each 3 µm distance covering both adherent cells and CCS were acquired every minute for a total of 4 hours.

Alternatively, HeLa cells stably expressing eGFP were cultured and infected in 8 well chambered coverslips (µ-Slide 8 Well, Ibidi). Live cell microscopy was performed at a Stellaris 8 Confocal Microscope (Leica Microsystems) equipped with a live cell chamber at 35°C. Z-stacks of 13 slices with each 2.5 µm distance covering both adherent cells and CCS were acquired every 5 minutes for a total of 4 hours.

##### Isolation and staining of CCS

CCS in the supernatant of *C. psittaci*-infected cell cultures were separated by centrifugation (5 min, 300 x *g*, RT). For live staining, pellet was mixed with the indicated staining solutions. The following staining solutions were used: 10 mM FM4-64 in DMSO (Sigma-Aldrich) diluted 1:1000 in infection medium; 25 mg mL^-1^ Hoechst 33342 (Sigma-Aldrich) diluted 1:2000 in infection medium; 5 mM SYTOX Green in DMSO (Thermo Fisher Scientific) diluted 1:2000 in infection medium; annexin V Alexa Fluor 568 (Thermo Fisher Scientific) diluted 1:10 in annexin binding buffer (PBS supplemented with 400 mg L^-1^ CaCl_2_). CCS were transferred into a 15 well chambered coverslip (µ-Slide 15 Well 3D, Ibidi) and either examined directly (FM4-64, Hoechst and SYTOX staining) or examined after 15 min of incubation at RT (annexin V staining) using an LSM 780 CLSM (Carl Zeiss). For immunofluorescence staining, pellet was mixed with 4% paraformaldehyde in PBS and transferred into a poly-L-lysine coated 8 well chambered coverslip (µ-Slide 8 Well, Ibidi). After 30 min of incubation at RT, staining was continued as described for immunofluorescence assays.

##### Antibodies used for immunofluorescence assay

Antibodies were used at the following concentrations: mouse anti-Hsp60 (chlamydial, cHsp60; 1:600, Cat. No. ALX-804-072, Enzo Life Sciences), rabbit anti-cleaved caspase-3 (1:400, Cat. No. 9661, Cell Signaling Technology), Alexa Fluor 488-coupled goat anti-rabbit IgG (1:100, Cat. No. 111-545-144, Dianova), and Cy3-coupled goat anti-mouse IgG (1:200, Cat. No. 115-165-146, Dianova).

##### Quantification of CCS

The number of CCS in *C. psittaci*-infected cell cultures was determined under different inhibitory and medium conditions. The following inhibitors were used: blebbistatin (Sigma-Aldrich, 50 µM), jasplakinolide (Santa Cruz, 1 µM), latrunculin B (Merck, 0.5 µM), nocodazole (Sigma-Aldrich, 30 µM), wiskostatin (Sigma-Aldrich, 50 µM), Z-VAD-FMK (R&D-Systems, 2 µM), and Z-DEVD-FMK (R&D-Systems, 2 µM). In addition, experiments with calcium- and FBS-free infection medium supplemented with the indicated concentrations of calcium chloride (Roth) were performed. At indicated time points, infection medium was changed to conditional medium and cultivation was continued for additional 4 h. To quantify the number of CCS, supernatant of the infected cell cultures was transferred to a new 12 well plate. The average number of CCS of at least five visual fields was determined and normalized to the infected cell area.

##### Determination of proteolytic DEVD cleaving activity

To determine proteolytic DEVD cleaving activity, cells were cultured and infected in 8 well chambered coverslips (µ-Slide 8 Well, Ibidi). Proteolytic DEVD cleaving activity was determined using the Incucyte Caspase-3/7 Red Dye for Apoptosis (Sartorius) as described in the manufacturer’s protocol. Briefly, Incucyte Caspase-3/7 Red Dye was diluted in infection medium to a final assay concentration of 833 nM and added to *C. psittaci*-infected cell cultures at indicated time points. After further cultivation (35°C, 5% CO_2_), samples were analyzed using an LSM 780 CLSM (Carl Zeiss).

##### Calcium imaging

Imaging of intracellular calcium levels and distribution was performed by either Rhod 3 staining or live cell imaging of the dual calcium reporter cell line HeLa ER-LAR-Geco G-Geco (Stelzner et al., 2020).

For Rhod 3 staining, cell culture and infection were performed in 8 well chambered coverslips (µ-Slide 8 Well, Ibidi). Rhod-3 staining was performed using the Rhod-3 Calcium Imaging Kit (Thermo Fisher Scientific) as described in the manufacturer’s protocol. In brief, cells were washed with PBS twice and 150 µL per well of freshly prepared Rhod-3 loading buffer (1x PowerLoad concentrate, 10 µM Rhod-3 AM, 250 µM Probenecid in infection medium or calcium- and FBS-free infection medium) were added. After incubation (60 min, 35°C, 5% CO_2_), cells were washed three times with PBS and 150 µL per well of incubation buffer (250 µM Probenecid in infection medium or calcium- and FBS-free infection medium) were added. After incubation (60 min, 35°C, 5% CO_2_), cells were washed with PBS and 150 µL per well of 25 mg mL^-1^ Hoechst 33342 (Sigma-Aldrich) diluted 1:2000 in infection medium were added for live cell microscopy. Live cell microscopy was performed at a Stellaris 8 Confocal Microscope (Leica Microsystems).

For live cell calcium imaging, the dual calcium reporter cell line HeLa ER-LAR-Geco G-Geco (Stelzner et al., 2020) was cultured and infected in 8 well chambered coverslips (µ-Slide 8 Well, Ibidi). This cell line stably expresses a red fluorescent endoplasmic reticulum (ER)-targeted fluorescent calcium sensor (ER-LAR-Geco) in addition to a green fluorescent cytosolic calcium indicator (G-Geco). For imaging of cytosolic calcium levels, the green channel was monitored using a Stellaris 8 Confocal Microscope (Leica Microsystems) equipped with a live cell chamber at 35°C. Z-stacks of 13 slices with each 2.5 µm distance covering both adherent cells and CCS were acquired every 2.5 minutes for a total of 4 hours.

